# No genetic evidence for involvement of alcohol dehydrogenase genes in risk for Parkinson’s disease

**DOI:** 10.1101/784405

**Authors:** Jonggeol Jeffrey Kim, Sara Bandres-Ciga, Cornelis Blauwendraat, International Parkinson’s Disease Genomics Consortium, Ziv Gan-Or

## Abstract

Multiple genes have been implicated in Parkinson’s disease (PD), including causal gene variants and risk variants typically identified using genome-wide association studies (GWAS). Variants in the alcohol dehydrogenase genes *ADH1C* and *ADH1B* are among the genes that have been associated with PD, suggesting that this family of genes may be important in PD. As part of the International Parkinson’s Disease Genomics Consortium’s (IPDGC) efforts to scrutinize previously reported risk factors for PD, we explored genetic variation in the alcohol dehydrogenase genes *ADH1A, ADH1B, ADH1C, ADH4, ADH5, ADH6*, and *ADH7* using imputed GWAS data from 15,097 cases and 17,337 healthy controls. Rare-variant association tests and single-variant score tests did not show any statistically significant association of alcohol dehydrogenase genetic variation with the risk for PD.

## Introduction

Parkinson’s disease (PD) is a progressive neurodegenerative disorder with a complex polygenic inheritance influenced by the interplay of genetic, aging and environmental factors. A variant in the gene encoding the Alcohol Dehydrogenase 1C (Class I) Gamma Polypeptide (*ADH1C*), an enzyme that catalyzes the oxidation of alcohol to acetaldehyde, was implicated as a potential risk factor involved in sporadic PD (Buervenich et al., 2005). Screening an international cohort of 1,076 PD patients of European ancestry and 940 matched controls, suggested that a rare nonsense variant (p.Gly78Ter) was significantly overrepresented in PD patients. Similarly, the variant p.His48Arg in the Alcohol Dehydrogenase 1B (Class I), Beta Polypeptide (*ADH1B*) was associated with PD in women in a cohort of 629 PD patients and 865 control participants(García-Martín et al., 2019). No additional studies on these genes have been performed, therefore the contribution of alcohol dehydrogenase genes to PD etiology has remained unclear. Here we investigate the potential association of common and rare variants in the alcohol dehydrogenase gene family with PD, in a large case-control series.

## Methods

To investigate the role of alcohol dehydrogenase genes and the effects of their genetic variation on the risk of PD, we utilized the International Parkinson’s Disease Genomics Consortium (IPDGC) genome-wide association study (GWAS) data consisting of 15,097 cases and 17,337 healthy controls (Supplementary Table 1) (Nalls et al., n.d.). The data underwent standard quality control as previously described (Nalls et al., n.d.) and were annotated using KGGSeq (Li et al., 2012).

We analyzed both the whole IPDGC cohort as well as two sex-stratified subcohorts. Variant frequencies were determined using PLINK 1.9 (Chang et al., 2015). Gene-based burden analyses SKAT, SKAT-O, and CMC were performed by using RVTESTS (Zhan et al., 2016) to assess the cumulative effect of multiple rare variants (minor allele frequency ≤ 0.03) on the risk for PD according to default parameters. Single-variant score test was performed using RVTESTS to assess the association between single variants and PD. All analyses were adjusted by 10 principal components to account for population stratification, dataset, age at onset for cases or examination for controls, and sex. All results were corrected for multiple testing by Benjamini–Hochberg False Discovery Rate (FDR) correction.

## Results

We identified 602 variants within the alcohol dehydrogenase gene loci, including 25 coding variants. Only two loss of function variants were identified: the previously reported p.Gly78Ter (rs283413) *ADH1C* variant, and the p.Tyr22Ter (rs3919370) of *ADH4*. The latter variant’s minor allele frequency (MAF = 0.29) did not meet the threshold for inclusion in the gene-level burden analysis tests for rare variants. No evidence for an association between alcohol dehydrogenase rare genetic variation and PD was detected when gene-based burden analysis was performed.

**TABLE 1.** Gene-based burden analysis of alcohol dehydrogenase variants and risk for Parkinson’s disease

| Gene | CHR | BP ranges | Cohort Sex | Number of Variants | P-value |  |  |
| --- | --- | --- | --- | --- | --- | --- | --- |
|  |  |  |  |  | CMC | SKAT | SKAT-O |
| ADH1A | 4 | 100197522 - 100212185 | All | 10 | 0.303 | 0.318 | 0.455 |
|  |  |  | Female | 10 | 0.307 | 0.574 | 0.675 |
|  |  |  | Male | 10 | 0.542 | 0.914 | 0.707 |
| ADH1B | 4 | 100227526 - 100242572 | All | 13 | 0.657 | 0.562 | 0.631 |
|  |  |  | Female | 13 | 0.968 | 0.385 | 0.596 |
|  |  |  | Male | 13 | 0.643 | 0.977 | 0.842 |
| ADH1C | 4 | 100257648 - 100273917 | All | 15 | 0.898 | 0.442 | 0.618 |
|  |  |  | Female | 15 | 0.619 | 0.276 | 0.454 |
|  |  |  | Male | 14 | 0.338 | 0.903 | 0.365 |
| ADH4 | 4 | 100044832 - 100065449 | All | 27 | 0.277 | 0.29 | 0.377 |
|  |  |  | Female | 27 | 0.232 | 0.703 | 0.793 |
|  |  |  | Male | 27 | 0.743 | 0.95 | 0.536 |
| ADH5 | 4 | 99992129 - 100009931 | All | 19 | 0.892 | 0.521 | 0.601 |
|  |  |  | Female | 19 | 0.657 | 0.703 | 0.862 |
|  |  |  | Male | 18 | 0.568 | 0.988 | 0.597 |
| ADH6 | 4 | 100125878 - 100140403,<br>100123794 - 100140403 | All | 5 | 0.619 | 0.356 | 0.566 |
|  |  |  | Female | 6 | 0.685 | 0.679 | 0.799 |
|  |  |  | Male | 5 | 0.754 | 0.864 | 0.891 |
| ADH7 | 4 | 100333417 - 100356667,<br>100333417 - 100356291 | All | 29 | 0.463 | 0.694 | 0.33 |
|  |  |  | Female | 29 | 0.88 | 0.818 | 0.835 |
|  |  |  | Male | 29 | 0.403 | 0.952 | 0.24 |

CHR, chromosome; BP, base pair; CMC, combined multivariate and collapsing; SKAT, sequence Kernel association tests; SKAT-O, optimized SKAT.

At variant-level, single-score test analyses found no evidence for an association between alcohol dehydrogenase variants and PD (FDR corrected p > 0.99) (Supplementary Table 1). The *ADH1C* p.Gly78Ter variant was previously reported with a frequency of 0.020 in PD patients, and a frequency of 0.006 in controls (Buervenich et al., 2005). However, in our cohort this variant was not associated with PD, as the allele frequency in patients was 0.016, while frequency in controls was 0.012 (p > 0.3, FDR corrected p > 0.99, Table 2). This is consistent with the frequency reported in gnomAD European (non-Finnish) origin population (0.012) (Karczewski et al., 2019).

**TABLE 2.**
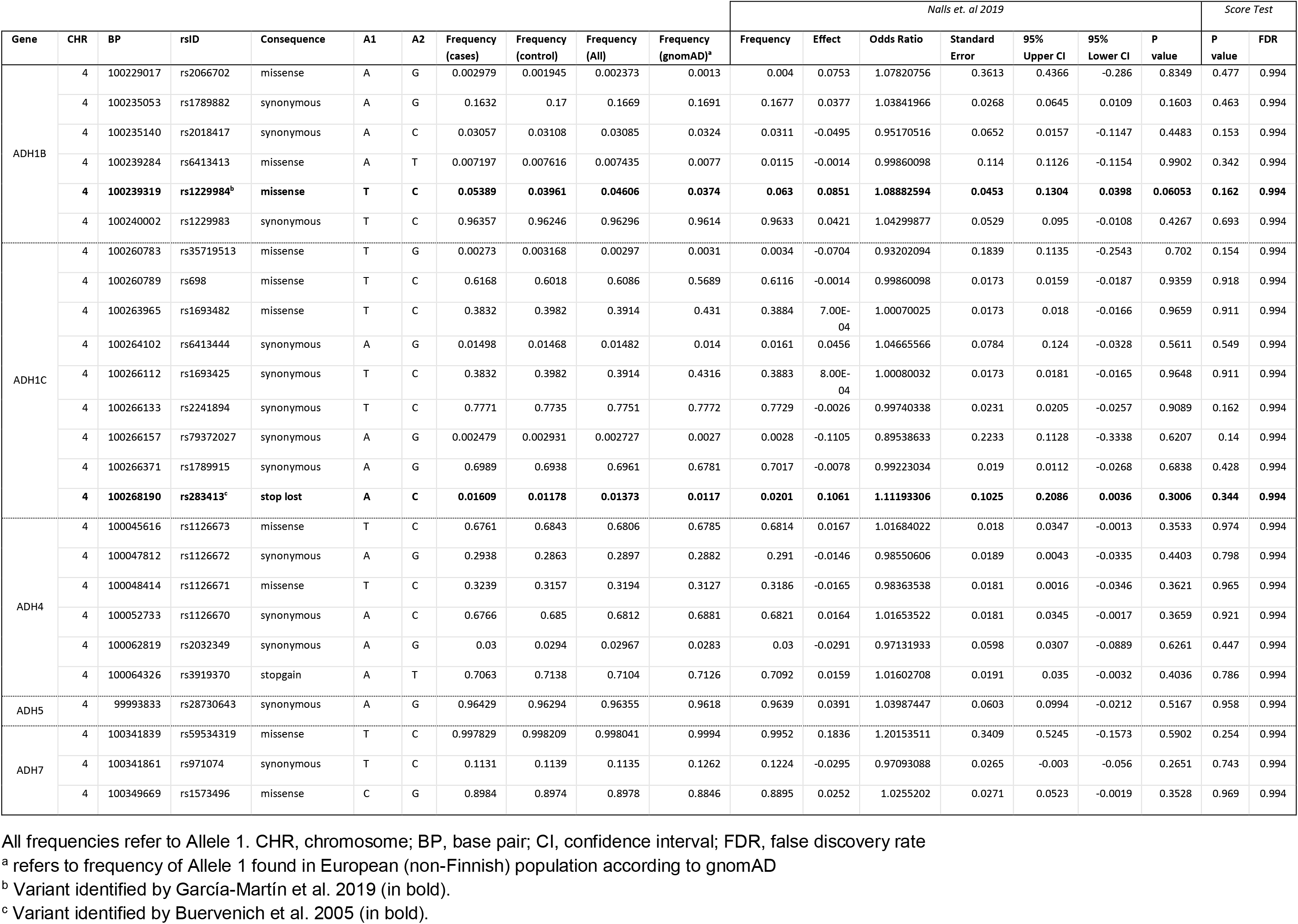
Coding variants of alcohol dehydrogenase genes and risk for Parkinson’s disease

| Gene | CHR | BP | rsID | Consequence | A1 | A2 | Frequency (cases) | Frequency (control) | Frequency (All) | Frequency (gnomAD)<sup>a</sup> | Nalls et. al 2019 |  |  |  |  |  |  | Score Test |  |
| --- | --- | --- | --- | --- | --- | --- | --- | --- | --- | --- | --- | --- | --- | --- | --- | --- | --- | --- | --- |
|  |  |  |  |  |  |  |  |  |  |  | Frequency | Effect | Odds Ratio | Standard Error | 95% Upper CI | 95% Lower CI | P value | P value | FDR |
| ADH1B | 4 | 100229017 | rs2066702 | missense | A | G | 0.002979 | 0.001945 | 0.002373 | 0.0013 | 0.004 | 0.0753 | 1.07820756 | 0.3613 | 0.4366 | -0.286 | 0.8349 | 0.477 | 0.994 |
|  | 4 | 100235053 | rs1789882 | synonymous | A | G | 0.1632 | 0.17 | 0.1669 | 0.1691 | 0.1677 | 0.0377 | 1.03841966 | 0.0268 | 0.0645 | 0.0109 | 0.1603 | 0.463 | 0.994 |
|  | 4 | 100235140 | rs2018417 | synonymous | A | C | 0.03057 | 0.03108 | 0.03085 | 0.0324 | 0.0311 | -0.0495 | 0.95170516 | 0.0652 | 0.0157 | -0.1147 | 0.4483 | 0.153 | 0.994 |
|  | 4 | 100239284 | rs6413413 | missense | A | T | 0.007197 | 0.007616 | 0.007435 | 0.0077 | 0.0115 | -0.0014 | 0.99860098 | 0.114 | 0.1126 | -0.1154 | 0.9902 | 0.342 | 0.994 |
|  | 4 | **100239319** | **rs1229984<sup>b</sup>** | **missense** | **T** | **C** | **0.05389** | **0.03961** | **0.04606** | **0.0374** | **0.063** | **0.0851** | **1.08882594** | **0.0453** | **0.1304** | **0.0398** | **0.06053** | **0.162** | **0.994** |
|  | 4 | 100240002 | rs1229983 | synonymous | T | C | 0.96357 | 0.96246 | 0.96296 | 0.9614 | 0.9633 | 0.0421 | 1.04299877 | 0.0529 | 0.095 | -0.0108 | 0.4267 | 0.693 | 0.994 |
| ADH1C | 4 | 100260783 | rs35719513 | missense | T | G | 0.00273 | 0.003168 | 0.00297 | 0.0031 | 0.0034 | -0.0704 | 0.93202094 | 0.1839 | 0.1135 | -0.2543 | 0.702 | 0.154 | 0.994 |
|  | 4 | 100260789 | rs698 | missense | T | C | 0.6168 | 0.6018 | 0.6086 | 0.5689 | 0.6116 | -0.0014 | 0.99860098 | 0.0173 | 0.0159 | -0.0187 | 0.9359 | 0.918 | 0.994 |
|  | 4 | 100263965 | rs1693482 | missense | T | C | 0.3832 | 0.3982 | 0.3914 | 0.431 | 0.3884 | 7.00E-04 | 1.00070025 | 0.0173 | 0.018 | -0.0166 | 0.9659 | 0.911 | 0.994 |
|  | 4 | 100264102 | rs6413444 | synonymous | A | G | 0.01498 | 0.01468 | 0.01482 | 0.014 | 0.0161 | 0.0456 | 1.04665566 | 0.0784 | 0.124 | -0.0328 | 0.5611 | 0.549 | 0.994 |
|  | 4 | 100266112 | rs1693425 | synonymous | T | C | 0.3832 | 0.3982 | 0.3914 | 0.4316 | 0.3883 | 8.00E-04 | 1.00080032 | 0.0173 | 0.0181 | -0.0165 | 0.9648 | 0.911 | 0.994 |
|  | 4 | 100266133 | rs2241894 | synonymous | T | C | 0.7771 | 0.7735 | 0.7751 | 0.7772 | 0.7729 | -0.0026 | 0.99740338 | 0.0231 | 0.0205 | -0.0257 | 0.9089 | 0.162 | 0.994 |
|  | 4 | 100266157 | rs79372027 | synonymous | A | G | 0.002479 | 0.002931 | 0.002727 | 0.0027 | 0.0028 | -0.1105 | 0.89538633 | 0.2233 | 0.1128 | -0.3338 | 0.6207 | 0.14 | 0.994 |
|  | 4 | 100266371 | rs1789915 | synonymous | A | G | 0.6989 | 0.6938 | 0.6961 | 0.6781 | 0.7017 | -0.0078 | 0.99223034 | 0.019 | 0.0112 | -0.0268 | 0.6838 | 0.428 | 0.994 |
|  | 4 | **100268190** | **rs283413<sup>c</sup>** | **stop lost** | **A** | **C** | **0.01609** | **0.01178** | **0.01373** | **0.0117** | **0.0201** | **0.1061** | **1.11193306** | **0.1025** | **0.2086** | **0.0036** | **0.3006** | **0.344** | **0.994** |
| ADH4 | 4 | 100045616 | rs1126673 | missense | T | C | 0.6761 | 0.6843 | 0.6806 | 0.6785 | 0.6814 | 0.0167 | 1.01684022 | 0.018 | 0.0347 | -0.0013 | 0.3533 | 0.974 | 0.994 |
|  | 4 | 100047812 | rs1126672 | synonymous | A | G | 0.2938 | 0.2863 | 0.2897 | 0.2882 | 0.291 | -0.0146 | 0.98550606 | 0.0189 | 0.0043 | -0.0335 | 0.4403 | 0.798 | 0.994 |
|  | 4 | 100048414 | rs1126671 | missense | T | C | 0.3239 | 0.3157 | 0.3194 | 0.3127 | 0.3186 | -0.0165 | 0.98363538 | 0.0181 | 0.0016 | -0.0346 | 0.3621 | 0.965 | 0.994 |
|  | 4 | 100052733 | rs1126670 | synonymous | A | C | 0.6766 | 0.685 | 0.6812 | 0.6881 | 0.6821 | 0.0164 | 1.01653522 | 0.0181 | 0.0345 | -0.0017 | 0.3659 | 0.921 | 0.994 |
|  | 4 | 100062819 | rs2032349 | synonymous | A | G | 0.03 | 0.0294 | 0.02967 | 0.0283 | 0.03 | -0.0291 | 0.97131933 | 0.0598 | 0.0307 | -0.0889 | 0.6261 | 0.447 | 0.994 |
|  | 4 | 100064326 | rs3919370 | stopgain | A | T | 0.7063 | 0.7138 | 0.7104 | 0.7126 | 0.7092 | 0.0159 | 1.01602708 | 0.0191 | 0.035 | -0.0032 | 0.4036 | 0.786 | 0.994 |
| ADH5 | 4 | 99993833 | rs28730643 | synonymous | A | G | 0.96429 | 0.96294 | 0.96355 | 0.9618 | 0.9639 | 0.0391 | 1.03987447 | 0.0603 | 0.0994 | -0.0212 | 0.5167 | 0.958 | 0.994 |
| ADH7 | 4 | 100341839 | rs59534319 | missense | T | C | 0.997829 | 0.998209 | 0.998041 | 0.9994 | 0.9952 | 0.1836 | 1.20153511 | 0.3409 | 0.5245 | -0.1573 | 0.5902 | 0.254 | 0.994 |
|  | 4 | 100341861 | rs971074 | synonymous | T | C | 0.1131 | 0.1139 | 0.1135 | 0.1262 | 0.1224 | -0.0295 | 0.97093088 | 0.0265 | -0.003 | -0.056 | 0.2651 | 0.743 | 0.994 |
|  | 4 | 100349669 | rs1573496 | missense | C | G | 0.8984 | 0.8974 | 0.8978 | 0.8846 | 0.8895 | 0.0252 | 1.0255202 | 0.0271 | 0.0523 | -0.0019 | 0.3528 | 0.969 | 0.994 |

All frequencies refer to Allele 1. CHR, chromosome; BP, base pair; CI, confidence interval; FDR, false discovery rate
<sup>a</sup> refers to frequency of Allele 1 found in European (non-Finnish) population according to gnomAD
<sup>b</sup> Variant identified by García-Martín et al. 2019 (in bold).
<sup>c</sup> Variant identified by Buervenich et al. 2005 (in bold).

Similarly, *ADH1B* p.His48Arg was previously reported to be associated with PD in women, with a frequency of 0.015 in PD patients and a frequency of 0.006 in controls (García-Martín et al., 2019). However, analysis of the IPDGC female subcohort showed no association with PD, as the effect allele frequency was 0.053 in patients and 0.040 in controls (P > 0.16, FDR corrected p > 0.89) (Table 2).

When summary statistics from 27,823 PD cases and 443,190 controls (Nalls et al., n.d.) were further assessed, no significant association was identified (Figure 1). These summary statistics included the cohort that was analyzed in this manuscript. Our results are based on the largest series of PD patients and controls to date with 100% statistical power to detect variants associated with PD at a MAF < 0.01, odds ratio = 1.5 and an α of 0.05.

**Figure 1.**
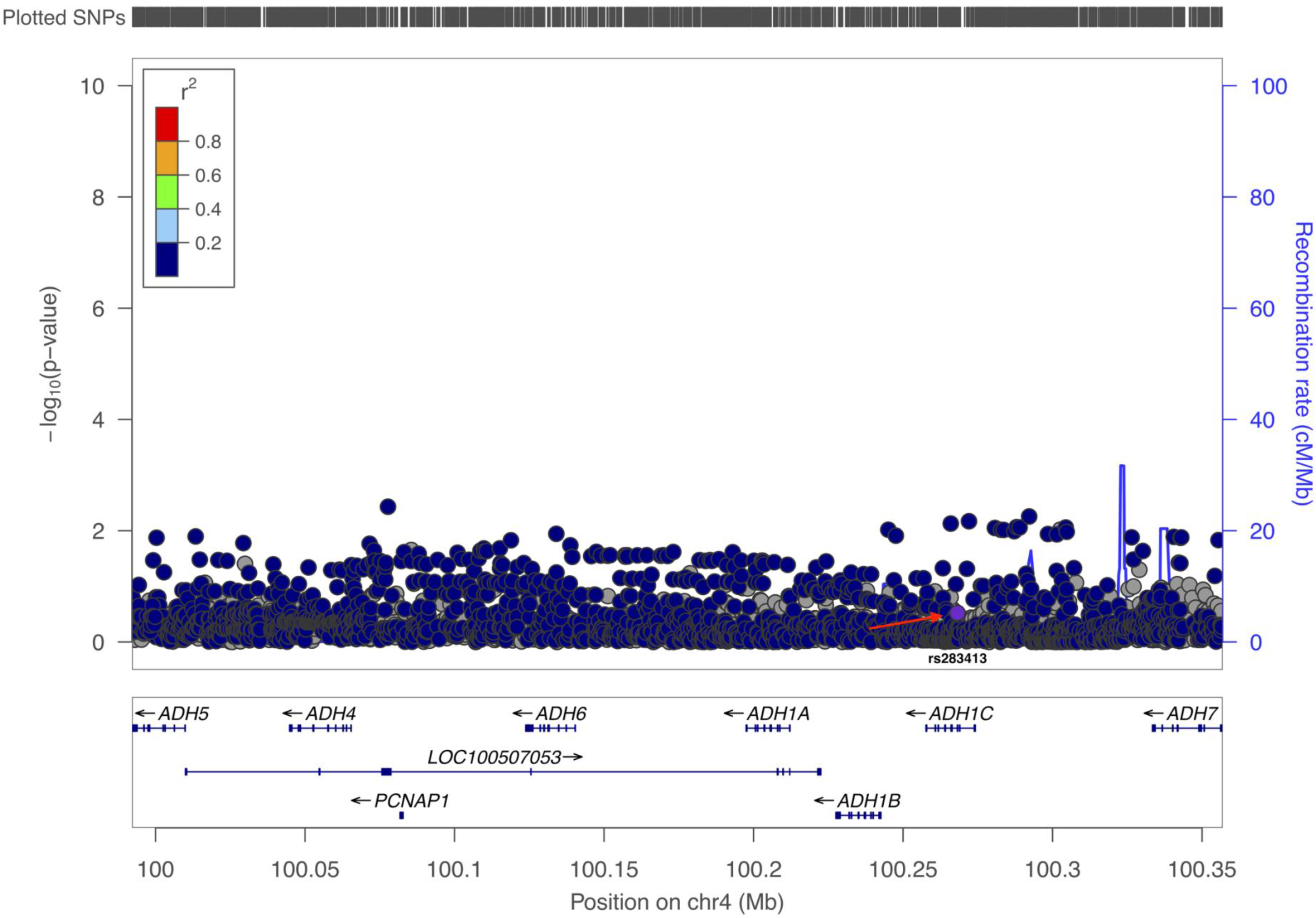
Locus zoom plot of the locus including the alcohol dehydrogenase genes, demonstrating lack of association with Parkinson’s disease risk. The violet circle with red arrow denotes the p.Gly78Ter variant of *ADH1C* (rs283413).

## Discussion

Our analyses, based on the largest PD genetic dataset to date, do not support the hypothesis that alcohol dehydrogenase genetic variants, including the *ADH1C* p.Gly78Ter stop loss variant and the *ADH1B* p.His48Arg missense variant, are risk or disease-causing factors for PD in the European population. The originally described variant, as well as other identified variants in these genes, have similar frequencies in patients and controls, and are therefore likely benign. A series of gene-based burden analyses and single variant analyses also failed to show a cumulative effect of rare variation in these genes on the risk for PD in the European population. In conclusion, our results do not support a role for common and rare variants in genes encoding the family of alcohol dehydrogenases as genetic risk factors or causal variants in PD. We suggest that these results should be taken into consideration when performing functional work.

## Acknowledgements

We would like to thank all of the subjects who donated their time and biological samples to be a part of this study. We also would like to thank all members of the International Parkinson Disease Genomics Consortium (IPDGC). See for a complete overview of members, acknowledgements and funding http://pdgenetics.org/partners. We would like to thank Sadie Zacharzuk for her assistance. The authors would like to thank the Genome Aggregation Database (gnomAD) and the groups that provided exome and genome variant data to this resource. A full list of contributing groups can be found at https://gnomad.broadinstitute.org/about.

