## Supplemental Information for "No genetic evidence for involvement of alcohol dehydrogenase genes in risk for Parkinson’s disease"

### Supplementary Information

**Supplementary Introduction**

Parkinson's disease (PD) is a progressive neurodegenerative disorder, pathologically characterized by degeneration of midbrain dopaminergic neurons and the presence of Lewy bodies. Key symptoms of PD are tremor, rigidity, bradykinesia, and gait instability. Other non-motor symptoms include constipation (Lin et al., 2014), hyposmia (Xiao et al., 2014), REM sleep behavior disorder (Sixel-Döring et al., 2016), autonomic dysfunction (De Pablo-Fernandez et al., 2017), depression, anxiety (Reijnders et al., 2008) (Aarsland et al., 2009) (Weintraub et al., 2015), and cognitive impairment (Miyasaki, n.d.). PD currently has no cure and treatments focus on minimizing symptoms and improving the quality of life of the patient. As the fastest growing neurological disorder in disability-adjusted life years, deaths, and prevalence (GBD 2016 Neurology Collaborators, 2019), PD represents a growing and significant burden on the global economy and society.

PD has a complex polygenic inheritance influenced by the interplay of genetic, aging and environmental factors (Billingsley et al., 2018). Age is the strongest PD risk factor and sex may also be a contributing factor; PD is disproportionately more prevalent in men (Van Den Eeden, 2003; Wright Willis et al., 2010). Numerous studies have shown that known and unknown environmental factors can contribute to PD risk. Pesticides, head injury, well water consumption (Noyce et al., 2012), and low-fat dairy consumption (Hughes et al., 2017) have been associated with increased risk for PD, while other factors such as tobacco usage, coffee, and alcohol have been associated with reduced risk for PD (Ascherio and Schwarzschild, 2016).

During the last two decades, large strides have been made in understanding the contribution of genetics to PD. Mutations/variants in multiple genes, including *SNCA, PRKN, LRRK2* and *GBA*, among others, are found in both familial and idiopathic PD cases (Reed et al., 2019). Multiple monogenic forms of PD have been discovered, but they only explain a fraction of PD with clear inheritance patterns. Current understanding of PD genetics have followed the common-disease-common-variant hypothesis, which postulates that multiple common variants with low risk and low penetrance in the general population can add up to cause complex but common diseases such as PD. The largest genome-wide association study of PD to date has identified 90 independent risk signals within 78 loci (Nalls et al., n.d.). However, these variants only explain 26-36% of the heritable risk of PD. There are multiple possible explanations for the “missing heritability problem”, including genetic structural variations and rare variants (Manolio et al., 2009).

Alcohol dehydrogenases (ADH) are a group of enzymes that oxidize alcohol to acetaldehyde and are crucial in the alcohol metabolism pathway (Cederbaum, 2012). The most common ADH is ADH (Class I), a dimer protein complex made of subunits encoded by the genes *ADH1A*, *ADH1B*, and *ADH1C*. A variant in the gene encoding the Alcohol Dehydrogenase 1C (Class I) Gamma Polypeptide (*ADH1C*) has been implicated as a potential risk factor for PD (Buervenich et al., 2005). A screening of an international cohort of 1,076 PD patients of European ancestry and 940 matched controls identified a rare nonsense variant (p.Gly78Ter, rs283413) which was significantly overrepresented in PD patients (odds ratio = 3.25, 95% CI = 1.31-8.05, P-value = 0.007). Subsequently, ADH1 and ADH1/4 knockout mice were produced as potential models for presymptomatic PD (Anvret et al., 2012). The knockout mice were reported to have significantly higher spontaneous locomotor activity compared to the wild-type mice, although there were no significant differences in olfactory function.

Similarly, the variant p.His48Arg (rs1229984) in the Alcohol Dehydrogenase 1B (Class I), Beta Polypeptide was associated with PD in female patients. Analyzing a cohort of 629 PD patients and 865 control participants, the authors found that this variant was significantly enriched in PD female patients (odds ratio = 2.13, 95% CI = 1.34–3.37, P-value < 0.001) (García-Martín et al., 2019) in a dominant model. The variant was not associated with PD age at onset or with PD in male patients.

Since then, the contribution of alcohol dehydrogenase genes to PD etiology has remained unclear. This is despite the increasing advancements in understanding the potential role of aldehyde dehydrogenase, another enzyme involved in alcohol metabolism, in PD and other neurodegenerative diseases (Grünblatt and Riederer, 2016; Wu et al., 2019). Here we investigate common and rare variants in the alcohol dehydrogenase gene family for their potential association with PD in a large case-control series. The effects of rare variants were collapsed to gene-level and analyzed using multiple different burden tests. Then both common and rare variants were analyzed using the single-variant score test.

#### Supplementary Methods

##### *Included data and quality control*

To investigate the role of alcohol dehydrogenase genes and their genetic variation on the risk for PD, we utilized the International Parkinson’s Disease Genomics Consortium (IPDGC) genome-wide association study (GWAS) data consisting of 15,097 PD patients and 17,337 healthy controls (Supplementary Table 1) (Nalls et al., n.d.). We analyzed both the entire IPDGC cohort, as well as sex-stratified subcohorts. The female subcohort consisted of 6,434 patients and 11,598 controls. The male subcohort consisted of 8,663 patients and 5,739 controls.

Sample inclusion criteria and quality control procedures have been previously described (Nalls et al., n.d.). In short, initial sample inclusion criteria included age at disease onset or last examination at 18 years of age or older, sample call rate of >95%, European ancestry confirmed through principal component analysis, no genetically ascertained relation to other samples in the meta-analysis at the cousin level or closer, and no heterozygosity outliers past +/- 15%. If a sample overlapped in two studies, it was removed from the larger study. All variants with missing call rate greater than 0.15 were removed using PLINK 1.9 (Chang et al., 2015). The remaining variants were annotated using KGGSeq (Li et al., 2012) for rsID, functional consequence, and Genome Aggregation Database (gnomAD) (Karczewski et al., 2019) frequencies.

*Statistical analysis*

Variant frequency was determined using PLINK 1.9. Gene-based burden analyses CMC, SKAT, and SKAT-O were performed by using RVTESTS (Zhan et al., 2016) to assess the cumulative effect of multiple rare variants (minor allele frequency ≤ 0.03) on the risk for PD according to default parameters. Single-variant score test was performed using RVTESTS to assess the association between single variants and PD.

CMC was chosen as it is a powerful burden analysis method that can effectively collapse rare variants at a gene-level, but works best under several assumptions: the relationships between the variants and PD have the same direction with similar magnitude, and the region being analyzed has a high proportion of causal variants compared to noncausal variants (Lee et al., 2014). To account for the chance that the above assumptions are not met, SKAT and SKAT-O were chosen as additional analysis methods. SKAT, a variance-component test, is statistically more powerful if the analyzed gene has more noncausal variants than causal and if the direction of the effect is more diverse. SKAT-O is a combination method that is more robust to the assumptions made by burden tests and variance-component tests, but is less powerful if either of the assumptions made by the burden test or variance-component test is true.

All analyses were adjusted by 10 principal components to account for population stratification, dataset, age at onset for patients or recruitment for controls, and sex. All results were corrected for multiple testing by Benjamini–Hochberg False Discovery Rate (FDR) correction.

**Complete Reference List**


Aarsland, D., Marsh, L., Schrag, A., 2009. Neuropsychiatric symptoms in Parkinson's disease. Mov. Disord. 24, 2175.

Anvret, A., Ran, C., Westerlund, M., Gellhaar, S., Lindqvist, E., Pernold, K., Lundströmer, K., Duester, G., Felder, M.R., Galter, D., Belin, A.C., 2012. Adh1 and Adh1/4 knockout mice as possible rodent models for presymptomatic Parkinson's disease. Behav. Brain Res. 227, 252–257.

Ascherio, A., Schwarzschild, M.A., 2016. The epidemiology of Parkinson's disease: risk factors and prevention. The Lancet Neurology. https://doi.org/10.1016/s1474-4422(16)30230-7

Billingsley, K.J., Bandres-Ciga, S., Saez-Atienzar, S., Singleton, A.B., 2018. Genetic risk factors in Parkinson's disease. Cell Tissue Res. 373, 9–20.

Buervenich, S., Carmine, A., Galter, D., Shahabi, H.N., Johnels, B., Holmberg, B., Ahlberg, J., Nissbrandt, H., Eerola, J., Hellström, O., Tienari, P.J., Matsuura, T., Ashizawa, T., Wüllner, U., Klockgether, T., Zimprich, A., Gasser, T., Hanson, M., Waseem, S., Singleton, A., McMahon, F.J., Anvret, M., Sydow, O., Olson, L., 2005. A Rare Truncating Mutation in ADH1C (G78Stop) Shows Significant Association With Parkinson Disease in a Large International Sample. Arch. Neurol. 62, 74.

Cederbaum, A.I., 2012. Alcohol metabolism. Clin. Liver Dis. 16, 667–685.

Chang, C.C., Chow, C.C., Tellier, L.C., Vattikuti, S., Purcell, S.M., Lee, J.J., 2015. Second-generation PLINK: rising to the challenge of larger and richer datasets. Gigascience 4, 7.

De Pablo-Fernandez, E., Tur, C., Revesz, T., Lees, A.J., Holton, J.L., Warner, T.T., 2017. Association of Autonomic Dysfunction With Disease Progression and Survival in Parkinson Disease. JAMA Neurol. 74, 970–976.

García-Martín, E., Diez-Fairen, M., Pastor, P., Gómez-Tabales, J., Alonso-Navarro, H., Alvarez, I., Cárcel, M., Aguilar, M., Agúndez, J.A.G., Jiménez-Jiménez, F.J., 2019. Association between the missense alcohol dehydrogenase rs1229984T variant with the risk for Parkinson's disease in women. Journal of Neurology. https://doi.org/10.1007/s00415-018-9136-9

GBD 2016 Neurology Collaborators, 2019. Global, regional, and national burden of neurological disorders, 1990-2016: a systematic analysis for the Global Burden of Disease Study 2016. Lancet Neurol. 18, 459–480.

Grünblatt, E., Riederer, P., 2016. Aldehyde dehydrogenase (ALDH) in Alzheimer's and Parkinson's disease. Journal of Neural Transmission. https://doi.org/10.1007/s00702-014-1320-1

Hughes, K.C., Gao, X., Kim, I.Y., Wang, M., Weisskopf, M.G., Schwarzschild, M.A., Ascherio, A., 2017. Intake of dairy foods and risk of Parkinson disease. Neurology 89, 46–52.

Karczewski, K.J., Francioli, L.C., Tiao, G., Cummings, B.B., Alföldi, J., Wang, Q., Collins, R.L., Laricchia, K.M., Ganna, A., Birnbaum, D.P., Gauthier, L.D., Brand, H., Solomonson, M., Watts, N.A., Rhodes, D., Singer-Berk, M., England, E.M., Seaby, E.G., Kosmicki, J.A., Walters, R.K., Tashman, K., Farjoun, Y., Banks, E., Poterba, T., Wang, A., Seed, C., Whiffin, N., Chong, J.X., Samocha, K.E., Pierce-Hoffman, E., Zappala, Z., O'Donnell-Luria, A.H., Minikel, E.V., Weisburd, B., Lek, M., Ware, J.S., Vittal, C., Armean, I.M., Bergelson, L., Cibulskis, K., Connolly, K.M., Covarrubias, M., Donnelly, S., Ferriera, S., Gabriel, S., Gentry, J., Gupta, N., Jeandet, T., Kaplan, D., Llanwarne, C., Munshi, R., Novod, S., Petrillo, N., Roazen, D., Ruano-Rubio, V., Saltzman, A., Schleicher, M., Soto, J., Tibbetts,

K., Tolonen, C., Wade, G., Talkowski, M.E., The Genome Aggregation Database Consortium, Neale, B.M., Daly, M.J., MacArthur, D.G., 2019. Variation across 141,456 human exomes and genomes reveals the spectrum of loss-of-function intolerance across human protein-coding genes. bioRxiv. https://doi.org/10.1101/531210
Lee, S., Abecasis, G.R., Boehnke, M., Lin, X., 2014. Rare-variant association analysis: study designs and statistical tests. Am. J. Hum. Genet. 95, 5–23.
Li, M.-X., Gui, H.-S., Kwan, J.S.H., Bao, S.-Y., Sham, P.C., 2012. A comprehensive framework for prioritizing variants in exome sequencing studies of Mendelian diseases. Nucleic Acids Res. 40, e53.
Lin, C.-H., Lin, J.-W., Liu, Y.-C., Chang, C.-H., Wu, R.-M., 2014. Risk of Parkinson's disease following severe constipation: a nationwide population-based cohort study. Parkinsonism Relat. Disord. 20, 1371–1375.
Manolio, T.A., Collins, F.S., Cox, N.J., Goldstein, D.B., Hindorff, L.A., Hunter, D.J., McCarthy, M.I., Ramos, E.M., Cardon, L.R., Chakravarti, A., Cho, J.H., Guttmacher, A.E., Kong, A., Kruglyak, L., Mardis, E., Rotimi, C.N., Slatkin, M., Valle, D., Whittemore, A.S., Boehnke, M., Clark, A.G., Eichler, E.E., Gibson, G., Haines, J.L., Mackay, T.F.C., McCarroll, S.A., Visscher, P.M., 2009. Finding the missing heritability of complex diseases. Nature 461, 747–753.
Miyasaki, J.M., n.d. Cognitive Decline in Parkinson Disease. Case Studies in Neuropalliative Care. https://doi.org/10.1017/9781108277365.022
Nalls, M.A., Blauwendraat, C., Vallerga, C.L., Heilbron, K., Bandres-Ciga, S., Chang, D., Tan, M., Kia, D.A., Noyce, A.J., Xue, A., Bras, J., Young, E., von Coelln, R., Simón-Sánchez, J., Schulte, C., Sharma, M., Krohn, L., Pihlstrom, L., Siitonen, A., Iwaki, H., Leonard, H., Faghri, F., Raphael Gibbs, J., Hernandez, D.G., Scholz, S.W., Botia, J.A., Martinez, M., Corvol, J.-C., Lesage, S., Jankovic, J., Shulman, L.M., Sutherland, M., Tienari, P., Majamaa, K., Toft, M., Andreassen, O.A., Bangale, T., Brice, A., Yang, J., Gan-Or, Z., Gasser, T., Heutink, P., Shulman, J.M., Wood, N., Hinds, D.A., Hardy, J.A., Morris, H.R., Gratten, J., Visscher, P.M., Graham, R.R., Singleton, A.B., The 23andMe Research Team, System Genomics of Parkinson's Disease (SGPD) Consortium, for the International Parkinson's Disease Genomics Consortium, n.d. Expanding Parkinson's disease genetics: novel risk loci, genomic context, causal insights and heritable risk. https://doi.org/10.1101/388165
Noyce, A.J., Bestwick, J.P., Silveira-Moriyama, L., Hawkes, C.H., Giovannoni, G., Lees, A.J., Schrag, A., 2012. Meta-analysis of early nonmotor features and risk factors for Parkinson disease. Ann. Neurol. 72, 893–901.
Reed, X., Bandrés-Ciga, S., Blauwendraat, C., Cookson, M.R., 2019. The role of monogenic genes in idiopathic Parkinson's disease. Neurobiology of Disease. https://doi.org/10.1016/j.nbd.2018.11.012
Reijnders, J.S.A.M., Ehrt, U., Weber, W.E.J., Aarsland, D., Leentjens, A.F.G., 2008. A systematic review of prevalence studies of depression in Parkinson's disease. Mov. Disord. 23, 183–9; quiz 313.
Sixel-Döring, F., Zimmermann, J., Wegener, A., Mollenhauer, B., Trenkwalder, C., 2016. The Evolution of REM Sleep Behavior Disorder in Early Parkinson Disease. Sleep 39, 1737–1742.
Van Den Eeden, S.K., 2003. Incidence of Parkinson's Disease: Variation by Age, Gender, and Race/Ethnicity. American Journal of Epidemiology. https://doi.org/10.1093/aje/kwg068
Weintraub, D., David, A.S., Evans, A.H., Grant, J.E., Stacy, M., 2015. Clinical spectrum of impulse control disorders in Parkinson's disease. Mov. Disord. 30, 121–127.
Wright Willis, A., Evanoff, B.A., Lian, M., Criswell, S.R., Racette, B.A., 2010. Geographic and ethnic variation in Parkinson disease: a population-based study of US Medicare beneficiaries. Neuroepidemiology 34, 143–151.

Wu, J., Kung, J., Dong, J., Chang, L., Xie, C., Habib, A., Hawes, S., Yang, N., Chen, V., Liu, Z., Evans, R., Liang, B., Sun, L., Ding, J., Yu, J., Saez-Atienzar, S., Tang, B., Khaliq, Z., Lin, D.-T., Le, W., Cai, H., 2019. Distinct Connectivity and Functionality of Aldehyde Dehydrogenase 1a1-Positive Nigrostriatal Dopaminergic Neurons in Motor Learning. Cell Rep. 28, 1167–1181.e7.

Xiao, Q., Chen, S., Le, W., 2014. Hyposmia: a possible biomarker of Parkinson's disease. Neurosci. Bull. 30, 134–140.

Zhan, X., Hu, Y., Li, B., Abecasis, G.R., Liu, D.J., 2016. RVTESTS: an efficient and comprehensive tool for rare variant association analysis using sequence data. Bioinformatics 32, 1423–1426.
